# SARS-CoV-2 S protein antagonizes type I interferon downstream signal pathway through interacting and attenuating phosphorylation of STAT1/STAT2

**DOI:** 10.1101/2022.06.06.494494

**Authors:** Wenjia Ni, Wenkang Li, Zeng Cai, Wenhua Guo, Yucheng Zheng, Yongliang Zhao, Zhixuan Wu, Simeng Liang, Jiajie Ye, Xiao Guo, Dan Zhou, Xiaoying Wu, Chanjuan Zhou, Mingliang Tang, Yu Chen, Ke Lan, Li Zhou, Ke Xu

## Abstract

Severe acute respiratory syndrome coronavirus 2 (SARS-CoV-2) may keep patients in a clinically asymptomatic state by blocking cellular innate antiviral immunity, but the molecular mechanism remains unclear. Here, we screened the viral proteins of SARS-CoV-2 and found that the spike (S) protein inhibits the activation of interferon-stimulated genes (ISGs) and even reduces the expression of these genes to below background values. Mechanistically, the S protein interacted with STAT1, STAT2, and IRF9 and impedes the phosphorylation of STAT1/STAT2, thus preventing the formation of the interferon-stimulating gene factor 3 (ISGF3) complex and inhibiting the downstream production of Interferon-stimulated genes (ISGs). Remarkably, we also have found that the inhibitory mechanism of the S protein was conservative among SARS-CoV-2 variants and other human coronaviruses, including SARS-CoV, MERS-CoV, HCoV-229E, HCoV-NL63, and HCoV-HKU1. Truncation studies indicated that the most conserved S2 domain played a major inhibitory role. Altogether, our findings unveil a new mechanism by which SARS-CoV-2 S protein attenuated the host’s antiviral immune response and provide new insights into the pathogenic mechanism of coronavirus.

## Introduction

Due to the high mutation rate and lack of efficient medicines, the pandemic caused by SARS-CoV-2 has become a great threat to global public health. Patients with COVID-19 mainly presented with symptoms such as fever, dry cough, and fatigue; however, some asymptomatic infections have also been observed (*1*). Clinical studies have reported that despite developing potent cytokines and chemokines in COVID-19 patients, the lack of interferon (IFN) response means that SARS-CoV-2 infection does not induce significant IFN production (*2, 3*).

Type I IFN is the first line of defense against viruses. Innate immune cells express pathogen recognition receptors (PRRs) to sense pathogen-associated molecular patterns (PAMP), including C-type lectin receptors (CLRs), NOD-like receptors (NLRs), RIG-I-like receptors (RLRs), and Toll-like receptors (TLRs) (*4, 5*). RNA viruses such as coronaviruses are recognized by cytoplasmic and endosomal RNA sensors, including RLRs and TLRs (TLR2, TLR7, and TLR8), respectively (*6-8*). It is demonstrated that the activation of TLR3 with the polyinosinic-polycytidylic acid (poly I: C) can inhibit coronavirus-related infection (*9*). Recognition of RNA viruses by TLRs and RLRs results in activation of transcription factors, such as nuclear factor-kappa light-chain enhancer (NF-κB) and interferon regulatory factor 3 (IRF3), leading to translocation to the nucleus and induction of pro-inflammatory cytokines, chemokines, and type I IFN expression (*10*). Afterward, IFN α/β activates the Janus kinase (JAK), signal transducer, and transcriptional activator (STAT) signaling pathway through the IFNAR. Upon JAK-STAT signaling, JAK1 and TYK2 mediate phosphorylation of STAT1/STAT2, thus forming an interferon-stimulating gene factor 3 (ISGF3) complex with(*11, 12*) IRF9. These complexes enter the nucleus to initiate the transcription of IFN-stimulated genes (ISGs) and subsequently the expression of antiviral proteins, including IFN-induced transmembrane proteins (IFITM) 1, 2, and 3, which restrict the infection of SARS-CoV-2 (*13, 14*).

Virus invasion triggers the host immune response, but the virus can also escape from the host immune response in various ways, such as inhibiting the production and secretion of IFN, blocking IFN signal transduction, etc. In response to host immune system attacks, structural and non-structural proteins of SARS-COV-2 (NSP1, NSP3, NSP6, NSP8, NSP12, NSP13, NSP14, ORF3a, ORF6, ORF7a, ORF7b, ORF8, ORF9b, M, and N proteins) have been reported to inhibit both types I IFN production and downstream signaling (*15-19*). Notably, all these proteins are intracellular expression proteins, but whether the membrane proteins of SARS-CoV-2, such as Spike (S) protein, also play a role is still unclear. The SARS-CoV-2 S protein mediates the viral membrane’s fusion with the host membrane and releases the viral RNA into host cells mainly through the Angiotensin-Converting Enzyme 2 (ACE2) (*20*). Thus, the S protein is the antigenic component of vaccine design (*21-23*). However, whether and how S protein antagonizes IFN downstream signaling is not clear.

This study investigated the interference between S protein and the host antiviral responses. We demonstrated that the S protein inhibits IFN downstream signaling transduction. On the mechanism, S protein impeded the phosphorylation of STAT1/STAT2 and interacted with STAT1, STAT2, and IRF9, thus preventing the formation of the interferon-stimulating gene factor 3 (ISGF3) complex. Finally, the S protein inhibited the downstream production of ISGs. Interestingly, the inhibitory mechanism of S protein was conservative in human coronavirus. In brief, our results provide evidence for the repression of innate immunity by S protein and provide a target for the treatment of COVID-19.

## Results

### SARS-CoV-2 Infection Inhibits ISG Expression in K18-hACE2 KI Mice

The virus infects cells and activates immune cells to express IFN and cytokines. However, it is reported that SARS-CoV-2 infection does not induce an efficient IFN-I response in COVID-19 patients, especially in critical patients (*24-26*). To explore the molecular mechanisms of this phenomenon, 12-18-week-old K18-hACE2 KI mice were intranasally (i.n.) inoculated with high-dose (3.0×10^5^ PFU) or low-dose (2.5×10^2^ PFU) of SARS-CoV-2 to mimic severe and mild viral infections, respectively (Fig. 1A). Subsequently, we extracted total mRNA from lung tissue of mice infected with SARS-CoV-2 at 2, 4, and 6 days post-infection (dpi), respectively. Then, we determined the mRNA levels of IFN-I-induced antiviral proteins (Ifitm1, Ifitm2, Ifitm3, Mx1, Isg15, Isg54, Isg56, and Ifng) in lung tissue of the K18-hACE2 KI mice by real-time quantitative PCR (qRT-PCR). We have found that the expression of IFN-γ and several ISGs was down-regulated in the lung tissues of both low-dose and high-dose virus-infected mice. In detail, for both the low-dose and high-dose groups, Ifitm1, Ifitm2, Mx1, Isg54, and Ifng were down-regulated to below background in late infection. Meanwhile, the expression of Ifitm2, Mx1, Isg15, Isg54, Isg56, and Ifng was down-regulated to lower as that time of viral infection extended. In addition, we also have found that Ifitm3, Isg15, and Isg56 activated at 2 dpi, but rapidly declined from 4 dpi to 6 dpi. Moreover, the activation of Ifitm3 and Isg15 in the high-dose virus-infected mice was significantly lower than that in the low-dose group at 2 dpi (Fig. 1B), suggesting that the high-dose virus infection caused more severe suppression of the innate immune system in mice. These data indicate that SARS-CoV-2 infection could effectively inhibit the expression of IFN-I-induced ISGs in K18-hACE2 KI mice.

**Figure 1.**
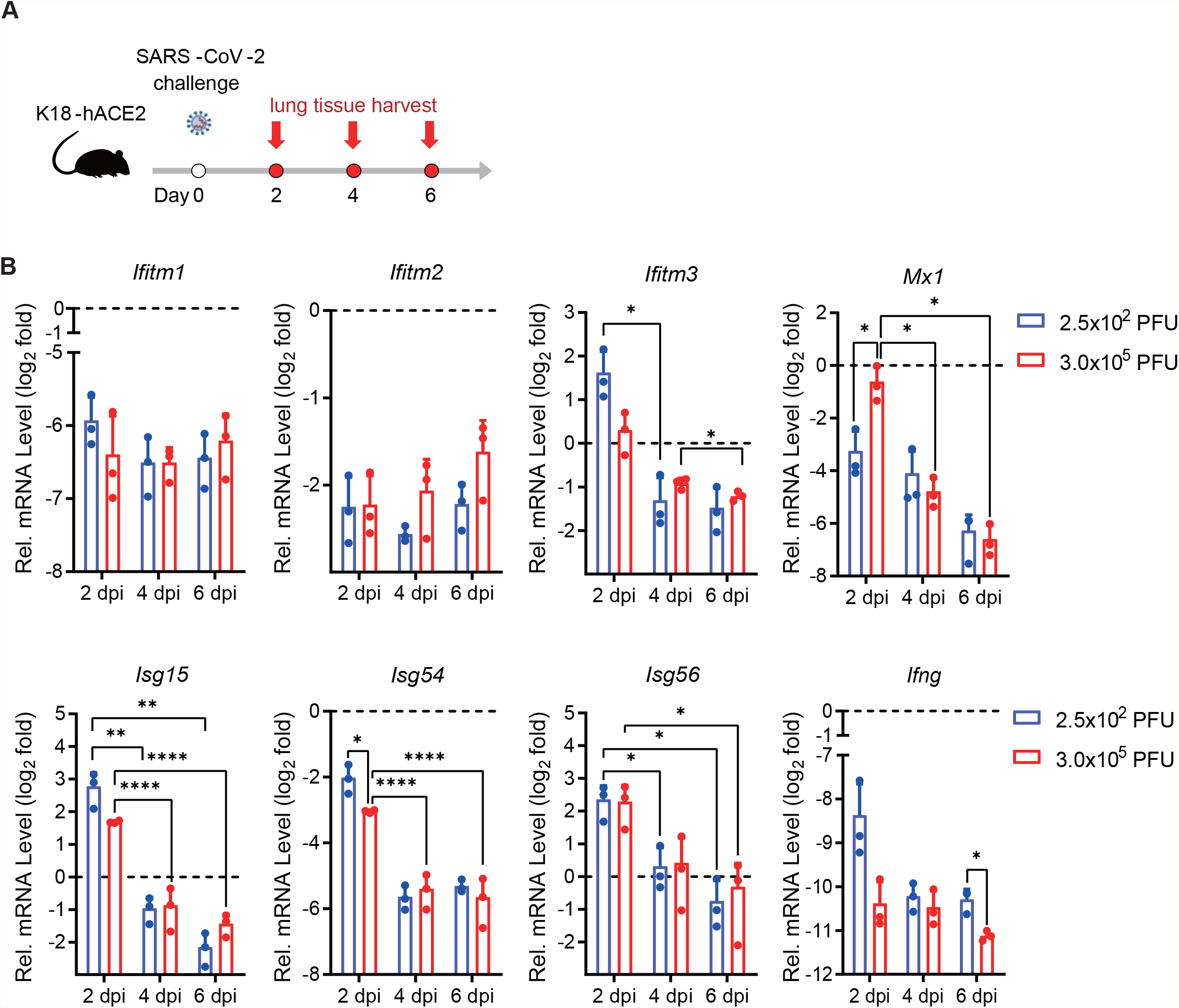
SARS-CoV-2 infection suppresses the mRNA level of antiviral proteins and cytokines in K18-hACE2 KI mice. (A) Schematic for SARS-CoV-2 infection. 12 to 18-week-old K18-hACE2 KI mice were anesthetized, followed by intranasal inoculation with 2.5×10^2^ or 3.0×10^5^ PFU SARS-CoV-2, respectively. Infected mice were euthanized at the indicated time. (B) mRNA levels of indicated genes were measured by relative quantification RT-PCR. The results were expressed as fold changes in gene RNA levels in infected mice relative to the uninfected control. Error bars represent mean ± SD from three independent experiments. Statistical significance was determined by Student’s t-test. *P < 0.05, **P < 0.01, ****P < 0.0001.

### SARS-CoV-2 Proteins Antagonize ISRE Activation and IFN-I-dependent ISG Induction

Since SARS-CoV-2 inhibited the activation of the IFN-I downstream signaling pathway, we suspected that the proteins of SARS-CoV-2 were involved in the regulation of the IFN-I signaling pathway. To test this hypothesis, we cloned 27 genes of SARS-CoV-2 after codon optimization and successfully expressed 19 proteins, including 11 non-structural proteins, 3 structural proteins, and 5 accessory proteins of SARS-CoV-2. We screened the genes by co-transfection of the plasmid encoding IFN-I-induced interferon-stimulated response element (ISRE) and the internal control plasmid pRL-SV40 into 293T cells, followed by the luciferase reporter assay. Besides, an NS1 expressing plasmid of influenza virus (A/WSN/33, H1N1) as a positive control (Fig. 2A). As expected, the positive control (NS1) and the NSP1, NSP3, NSP13, ORF3a, ORF7a, ORF7b, ORF8, M, and N proteins have been reported in other articles about their inhibitory mechanisms. Surprisingly, we also found that S, NSP5, NSP7, NSP10, NSP16 and ORF9b proteins significantly inhibited activation of the ISRE stimulated by IFN-I. We considered that the S protein acts as a structural protein to mediate viral invasion. Whereas whether S affects host innate immunity is still unknown, we first focused on the role of S protein in inhibiting ISRE activity. A dose-dependent assay revealed that the S protein inhibited IFN-I-induced activation of ISRE, IFITM3, and MxA promoter (Fig. 2B-D). We next overexpressed the S protein and examined the endogenous expressions of IFITM1, IFITM2, IFITM3, MxA, ISG15, ISG54, ISG56, and IFNγ induced by IFN-I through qRT-PCR analysis. We observed that those genes were significantly inhibited ∼200, 192, 8, 20, 115, 13, 72, and 60 folds, respectively (Fig. 2E). These results indicated that the S protein plays an important role in suppressing the host IFN response pathway.

**Figure 2.**
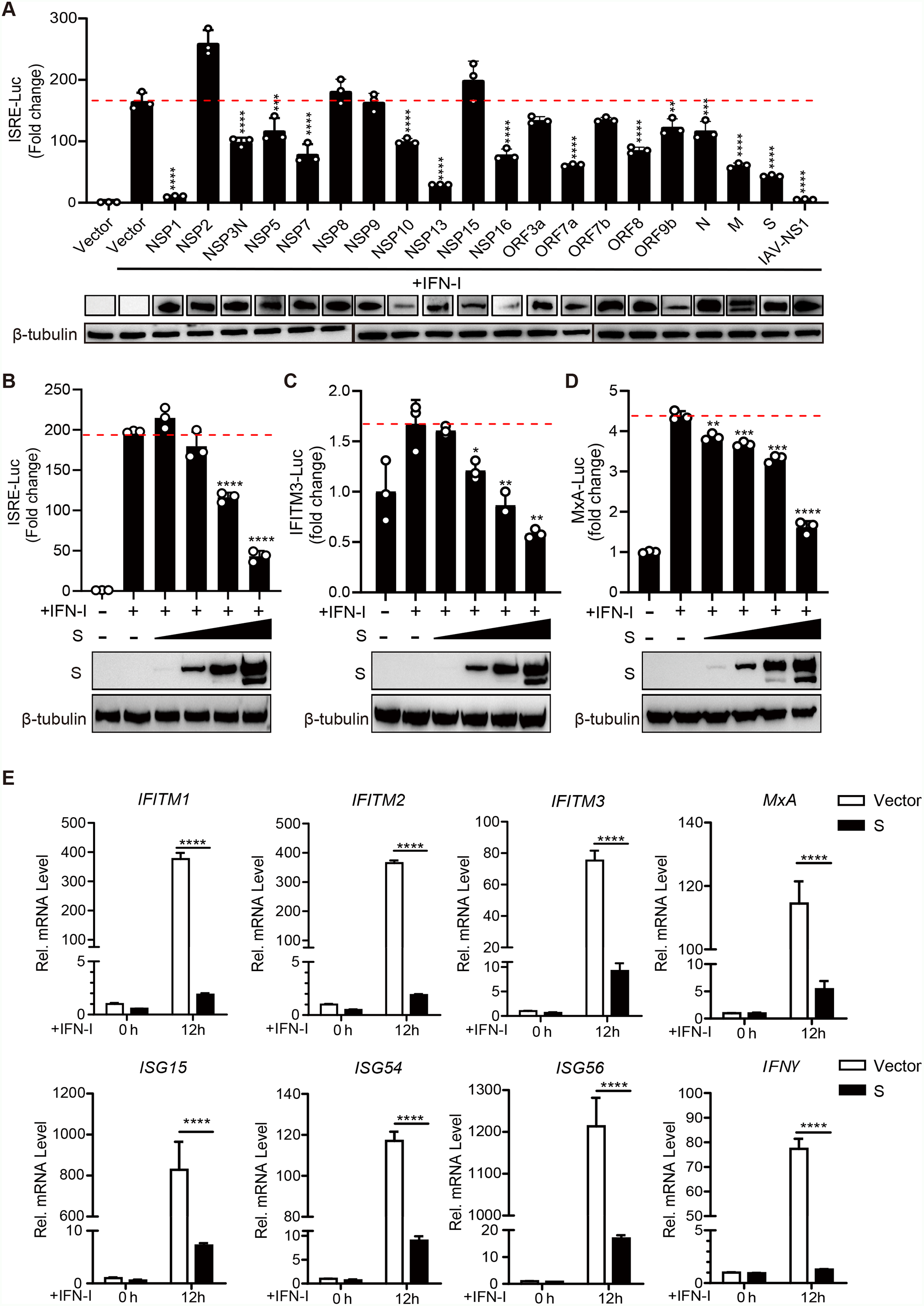
SARS-CoV-2 S proteins inhibit the activation of ISRE and the induction of ISGs. (A) 293T cells were co-transfected with pISRE-Luc, Renilla luciferase control plasmid pRL-SV40, and SARS-CoV-2 protein-expressing plasmid. 24 h after the initial transfection, the cells were stimulated with IFN-I (1,000 U/mL) for 12 h, and the luciferase assays were measured. (B-D) 293T cells were co-transfected with pISRE-Luc (B), pIFITM3-Luc (C), or pMxA-Luc (D) together with the pRL-SV40 plasmid, and the plasmid pCAGGS-Flag-S of 0, 0.01, 0.05, 0.25, and 1.25 μg. All promoter activity was measured upon IFN-I (1,000 U/mL) stimulated for 12 h. Error bars represent mean ± SD from three independent experiments. Statistical significance was determined by One-Way ANOVA test. *P < 0.05, **P < 0.01, ***P < 0.001, ****P < 0.0001. (E) HeLa cells were transfected with empty vector or SARS-CoV-2 S protein-expressing plasmids for 24 h and then stimulated with IFN-I (1,000 U/mL) for 12 h. mRNA expression levels of IFITM1, IFITM2, IFITM3, MxA, ISG15, ISG54, ISG56, and IFNγ in the collected cells were detected by qRT-PCR. Error bars represent mean ± SD from three independent experiments. Statistical significance was determined by Student’s t-test. ****P < 0.0001.

### SARS-CoV-2 S Protein Antagonizes the JAK-STAT Signal Pathway

The IFN-I receptor recognizes the secreted IFN-I, which is then transduced through the JAK-STAT signaling pathway and activates the expression of hundreds of ISGs. To investigate the mechanisms by which SARS-CoV-2 S protein antagonizes IFN-I downstream signaling pathway, we examined the phosphorylation of the STAT1/STAT2 within the JAK/STAT pathway in response to the IFN-I stimulation. Thus, HeLa cells were transiently expressing S protein or empty vector plasmid and followed by treatment with IFN-I for 6 h or 12 h, and then monitored expression and phosphorylation levels of STAT1/STAT2 by Western blot. As expected, IFN-I treatment lead to the phosphorylation of both STAT1/STAT2. Interestingly, overexpression of the SARS-CoV-2 S protein did not affect the expression levels of STAT1/STAT2 but reduced phosphorylation levels of STAT1/STAT2. And the decrease of p-STAT1/p-STAT2 at 12 hours was more than at 6 hours after IFN-I induction (Fig. 3A). Once STAT1/STAT2 are phosphorylated by IFN-I stimulation, pSTAT1 and pSTAT2 form a heterodimer, which recruits IRF9 to form the STAT1/STAT2/IRF9 complex (ISGF3), then ISGF3 translocates to the nucleus and binds to ISRE, thereby leads to the expression of ISGs. Next, we tested the effect of SARS-CoV-2 S protein on STAT1 nuclear translocation stimulated with IFN-I. HeLa cells were transfected with an S-expressing plasmid followed by treatment with IFN-I for 30 min and then analyzed by immunofluorescence microscopy. Immunofluorescence analysis showed that STAT1 dispersed in the cytoplasm with neither S expression nor IFN-I treatment. Subsequently, STAT1 was translocated to the nucleus under IFN-I treatment. However, in the presence of S protein, STAT1 remained in the cytoplasm even under IFN-I stimulation. (Fig. 3B). These results suggest that S protein inhibited the phosphorylation of STAT1/STAT2 and suppressed STAT1 nuclear translocation induced by IFN-I.

**Figure 3.**
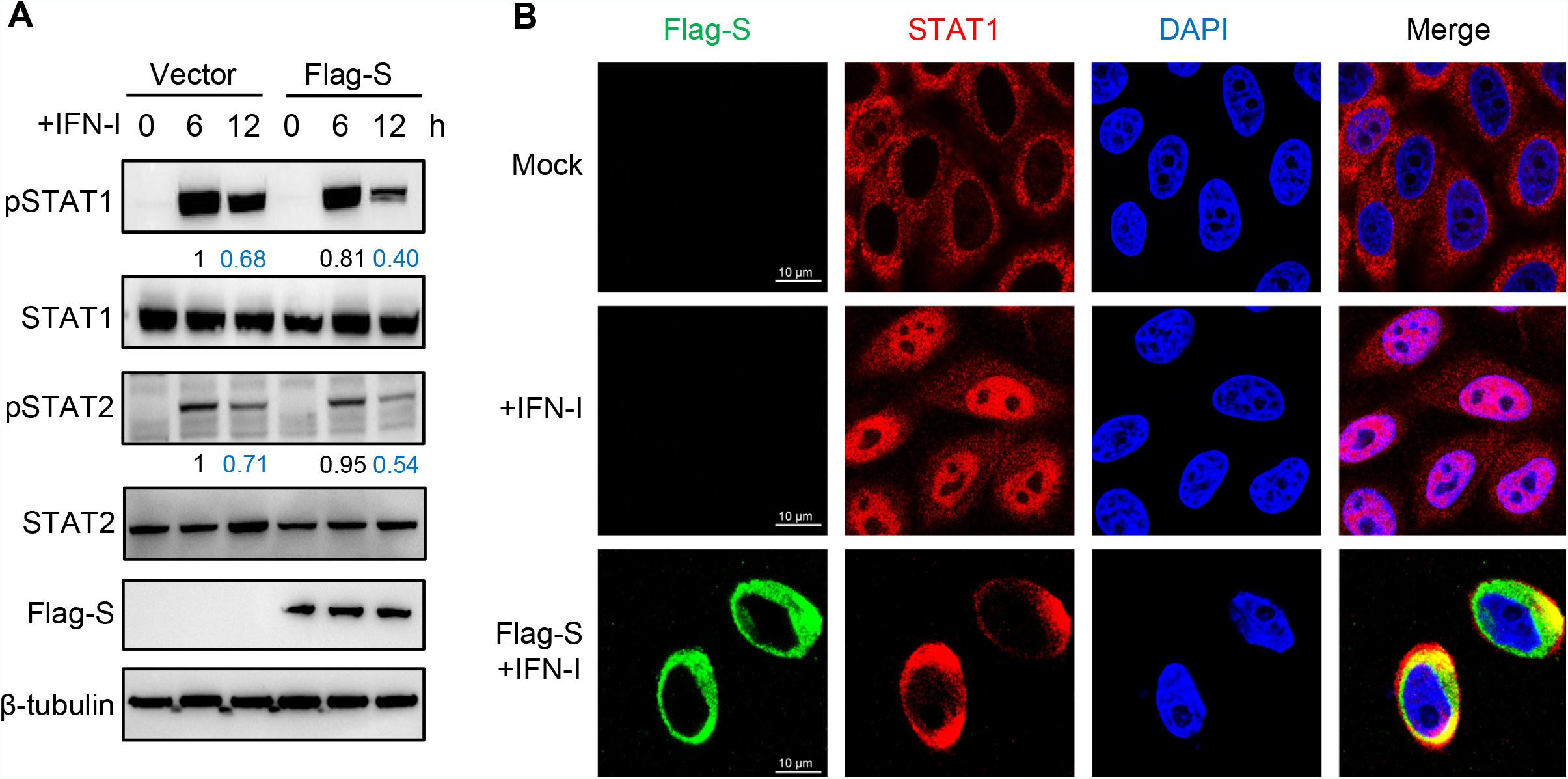
SARS-CoV-2 S protein inhibits phosphorylation of STAT1/STAT2 and blocks STAT1 nuclear translocation. (A) 293T cells were transfected with empty vector plasmid or SARS-CoV-2 S protein-expressing plasmids for 24 h and then stimulated with IFN-I for 6 and 12 h, and analyzed by Western blot using anti-pSTAT1, anti-total STAT1, anti-pSTAT2, and anti-total STAT2 antibodies. Protein band intensity was quantitated using ImageJ software. Data represent the means of three independent experiments. (B) HeLa cells were transfected with plasmid expression Flag-S for 24 h, and treated with IFN-I (1,000 U/mL) for 30 min. Cells were fixed and permeabilized, staining with anti-STAT1 and anti-FLAG as primary antibodies and anti-Alexa Fluor 488 and anti-Alexa Fluor 568 as secondary antibodies. Scale bar, 10 μm.

### SARS-CoV-2 S Directly Interacts with the ISGF3 Complex

Because the S protein inhibits the phosphorylation of STAT1/STAT2 and suppresses STAT1 nuclear translocation, we next examined whether S protein can interact with STAT1, STAT2, and IRF9, the component of ISGF3. We co-transfected the plasmid encoding S protein and STAT1, STAT2, and IRF9, respectively, followed by a co-immunoprecipitated (co-IP) assay. As shown in Fig. 4A, C, E, Flag-tagged S protein could precipitate with HA-tagged STAT1, STAT2, and IRF9. In addition, we overexpressed Flag-tagged S along with HA-tagged STAT1, STAT2, or IRF9 in HeLa cells, and examined their subcellular locations via immunofluorescence confocal microscopy. Our data showed that S protein obviously co-localized with STAT1, STAT2, and IRF9. Moreover, these colocalizations were located on the endoplasmic reticulum / Golgi apparatus in the cytoplasm (Fig. 4B, D, F). Next, we tested whether S protein can disturb the formation of the ISGF3 complex. We co-transfected the plasmid expressing STAT1, STAT2, and IRF9 with S protein, followed by a co-IP assay. The result showed that IRF9 increased binding to STAT1 but not STAT2 with IFN-I treatment. In contrast, IRF9 showed decreased binding to S protein and increased binding to STAT1 under IFN-I treatment (Fig. 4G). That’s to say, S protein competed against STAT1 for binding with IRF9 and STAT2. Together, our findings indicated that the SARS-CoV-2 S protein disturbs the formation of the ISGF3 complex, which in turn suppresses the JAK/STAT signal pathway.

**Figure 4.**
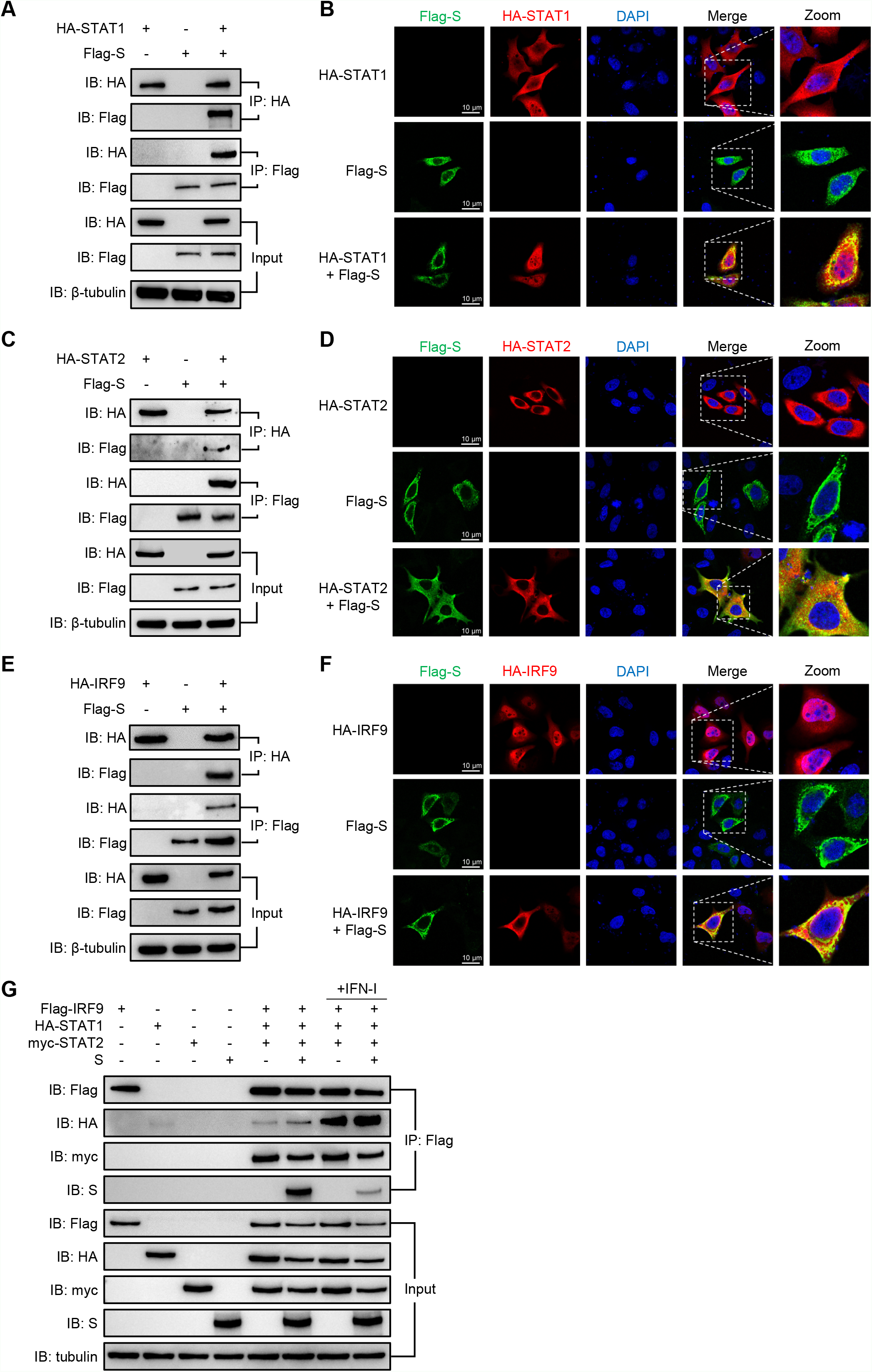
SARS-CoV-2 S directly interacts with ISGF3. (A, C, E) 293T cells were transfected with the plasmid as indicated. At 48 hpt, the cells were harvested and subjected to a Co-IP assay. The expression of HA-STAT1, HA-STAT2, HA-IRF9, Flag-S, and β-tubulin were shown as a loading control (Input). (B, D, F) HeLa cells were transfected with plasmid expression HA-STAT1, HA-STAT2, or HA-IRF9 along with Flag-S for 24 h. Cells were fixed and permeabilized, staining with anti-HA and anti-FLAG as primary antibodies, anti-Alexa Fluor 488 and anti-Alexa Fluor 568 as secondary antibodies. Scale bar, 10 μm. (G) 293T cells were transfected with the plasmid as indicated for 48 h, and then treated with IFN-I (1,000 U/mL) for 1 h. Cells were harvested and subjected to a Co-IP assay. The expression levels of Flag-IRF9, HA-STAT1, myc-STAT2, S, and β-tubulin were shown as a loading control (Input).

### The S2 Domain Is Mainly Responsible for the Inhibition of ISRE

To determine the pivotal domain of S protein on inhibitory effect on the JAK/STAT signal pathway, we split the full-length S protein into four domains, NTD (aa 12-306), RBD (aa 328-533), SD1/2 (aa 535-680), and S2 (aa 688-1273). 293T cells were transfected with the plasmids expressing the four domains, respectively, along with pISRE-Luc and pRL-SV40, followed by a dual-luciferase assay. The result showed that the S2 domain inhibited the activity of ISRE by 60%, distributing the major inhibitory effect. However, NTD showed no inhibitory effect. RBD and SD1/2 only slightly inhibited the activity of ISRE by ∼34% and 27%, respectively (Fig. 5A). To validate whether the S2 domain is the major suppressor of S protein, we test the effect of the S2 domain on STAT1 nuclear translocation stimulated with IFN-I as mentioned above. As shown in Fig. 5B, S2 domain suppressed STAT1 nuclear translocation induced by IFN-I. Moreover, we verified the S2 domain could interact with STAT1, STAT2, and IRF9 (Fig. 5C, E, G), and the immunofluorescence results confirmed these results (Fig. 5D, F, H). In conclusion, the S2 domain is mainly responsible for inhibiting the activity of ISRE.

**Figure 5.**
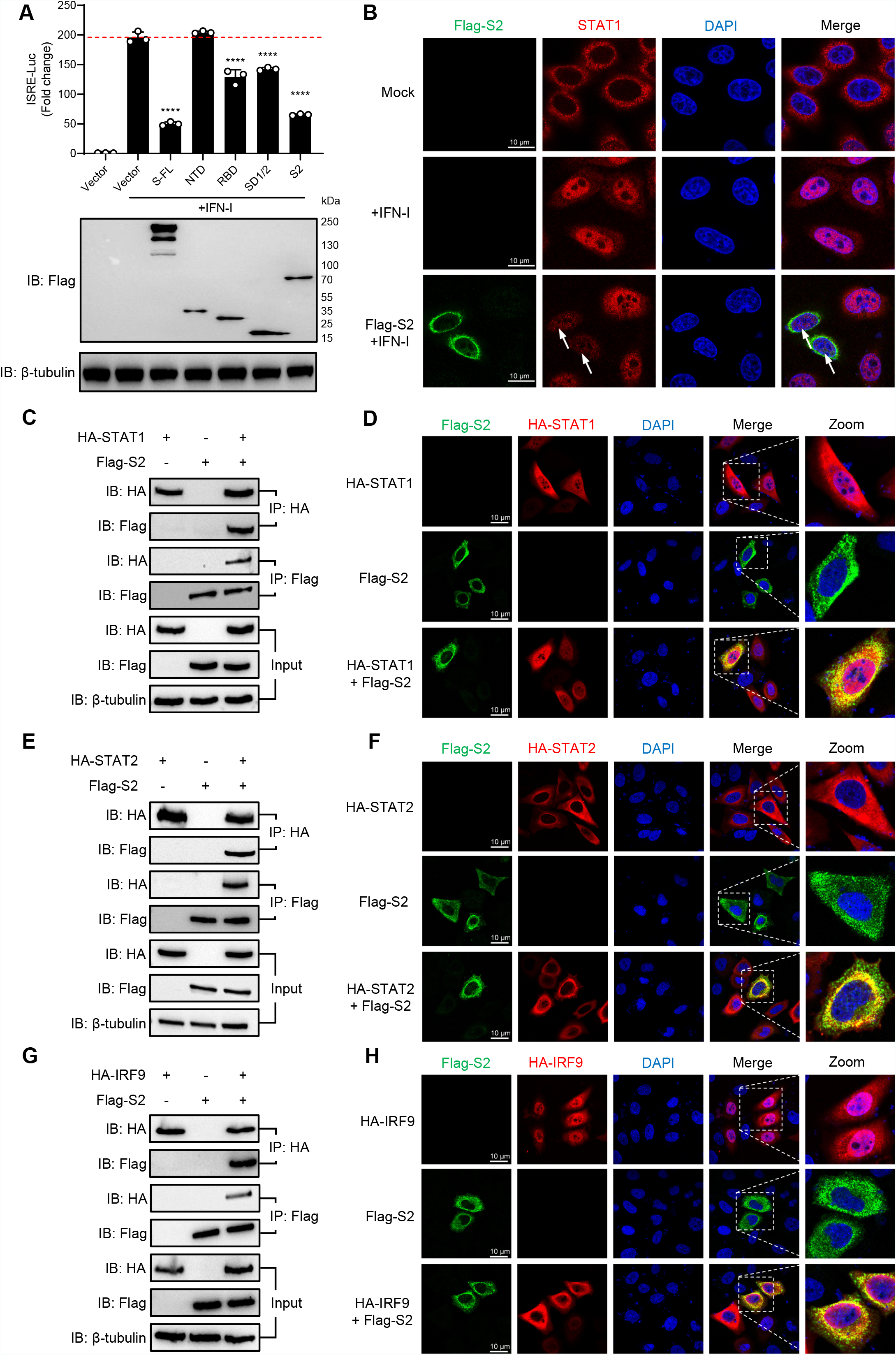
The S2 domain is mainly responsible for the inhibitioy mechanism. (A) 293T cells grown in 24-well plates were co-transfection with pISRE-Luc, pRL-SV40 control plasmid, and a plasmid expressing different S domain protein. At 24 hpt, cells were treated with IFN-I (1,000 U/mL) for 12 h, and the luciferase activity was measured. FL, full-length. Error bars represent mean ± SD from three independent experiments. Statistical significance was determined by One-Way ANOVA test, ****p < 0.0001. (B) HeLa cells were transfected with plasmid expression Flag-S2 for 24 h, treated with IFN-I (1,000 U/mL) for 30 min. Cells were fixed and permeabilized, staining with anti-STAT1 and anti-FLAG as primary antibodies, anti-Alexa Fluor 488 and anti-Alexa Fluor 568 as secondary antibodies. Scale bar, 10 μm. (C, E, G) 293T cells were transfected with the plasmid as indicated. At 48 hpt, the cells were harvested and subjected to a Co-IP assay. The expression of HA-STAT1, HA-STAT2, HA-IRF9, Flag-S2, and β-tubulin were shown as a loading control (Input). (D, F, H) HeLa cells were transfected with plasmid expression HA-STAT1, HA-STAT2, or HA-IRF9 along with Flag-S2 for 24 h. Cells were fixed and permeabilized, staining with anti-HA and anti-FLAG as primary antibodies and anti-Alexa Fluor 488 and anti-Alexa Fluor 568 as secondary antibodies. Scale bar, 10 μm.

### SARS-CoV-2 Variants and Other Human Coronavirus S Proteins Inhibit the Activity of ISRE

Since the S2 domain is conserved in SARS-CoV-2 variants S protein, we tested the inhibitory ability of prevalent SARA-CoV-2 variants on the activity of ISRE. We have found that the S proteins of the D614G variant, Alpha variant (B.1.1.7), Beta variant (B.1.351), Gamma variant (P.1), Delta variant (B.1.617.2), and Omicron variant (B.1.1.529) significantly inhibited the IFN-I-mediated ISRE activation, and the inhibitory ability had no significant difference from the wild type (Fig. 6A). Moreover, we aligned different human coronavirus S protein sequences, including SARS-CoV-2, SARS-CoV, MERS-CoV, HCoV-229E, HCoV-NL63, and HCoV-HKU1, and analyzed by Simplot software. It was shown that the S2 domain had a high similarity in different human coronavirus (Fig. 6C). The analysis result indicates that the inhibitory ability may be conservative in coronavirus. We next repeated the ISRE-luciferase assay to verify the hypothesis using plasmids expressing S2 domains of other human coronavirus S proteins. As expected, all tested coronavirus S proteins and S2 domains exerted a similar effect on ISRE (Fig. 6B and D). Our data implied that the S2 domain is mainly responsible for the inhibition of the activity of ISRE and that the inhibition of the activity of ISRE by coronavirus is ubiquitous.

**Figure 6.**
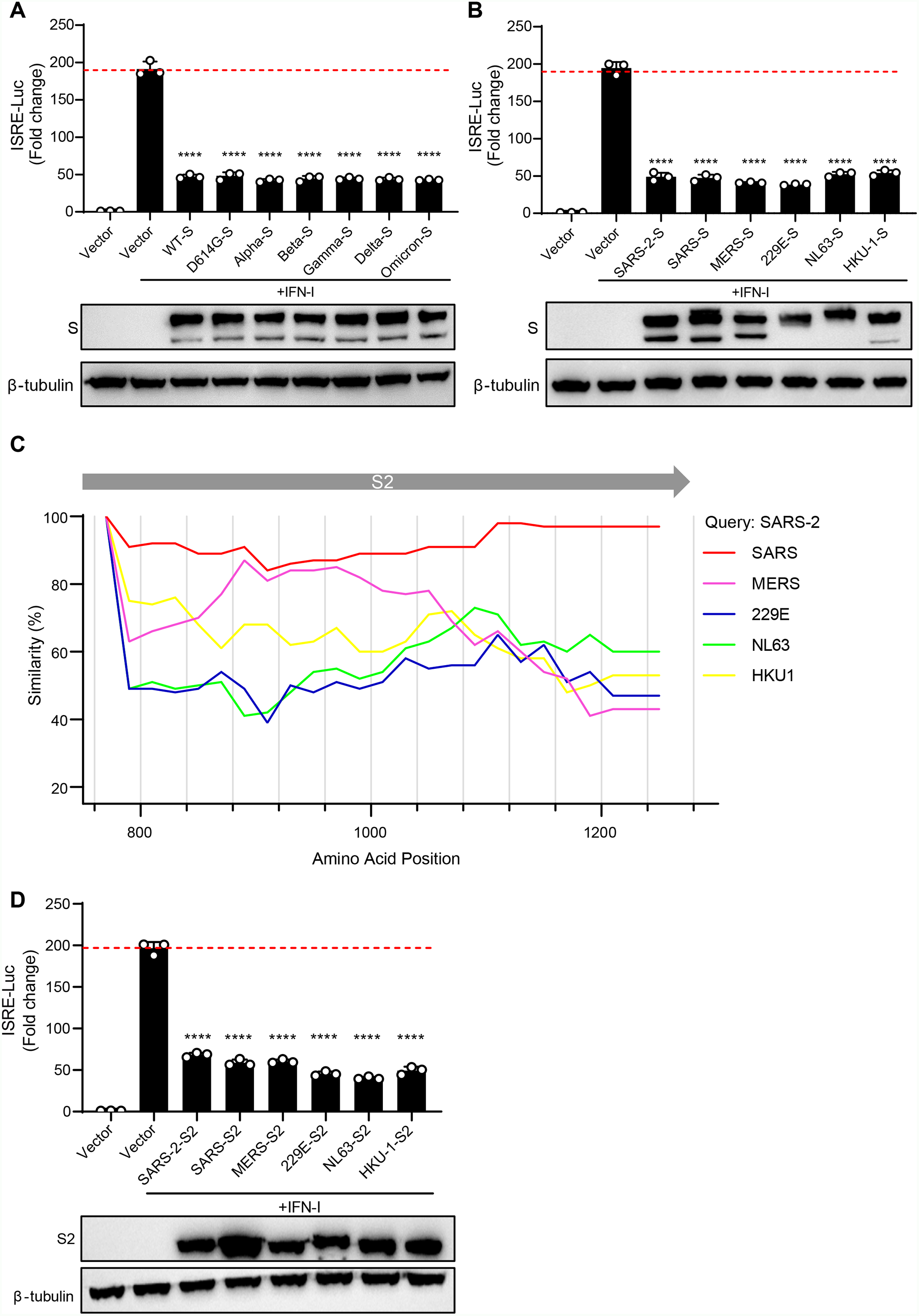
Comparison of IFN-I inhibition by different coronavirus S proteins. (A) 293T cells grown in 24-well plates were co-transfection with pISRE-Luc, pRL-SV40 control plasmid, and a plasmid expressing different SARS-CoV-2 variants S proteins. At 24 hpt, cells were treated with IFN-I (1,000 U/mL) for 12 h, and the luciferase activity was measured. (B) 293T cells grown in 24-well plates were co-transfection with pISRE-Luc, pRL-SV40 control plasmid, and a plasmid expressing different coronavirus S proteins. At 24 hpt, cells were treated with IFN-I (1,000 U/mL) for 12 h, and the luciferase activity was measured. (C) Simplot analyses the similarity of different human coronavirus S protein sequences. The abscissa represents the amino acid position of the SARS-CoV-2 S protein. (D) 293T cells grown in 24-well plates were co-transfection with pISRE-Luc, pRL-SV40 control plasmid, and a plasmid expressing different coronavirus S2 domains. At 24 hpt, cells were treated with IFN-I (1,000 U/mL) for 12 h, and the luciferase activity was measured. Error bars represent mean ± SD from three independent experiments. Statistical significance was determined by One-Way ANOVA test, ****p < 0.0001.

**Figure 7.**
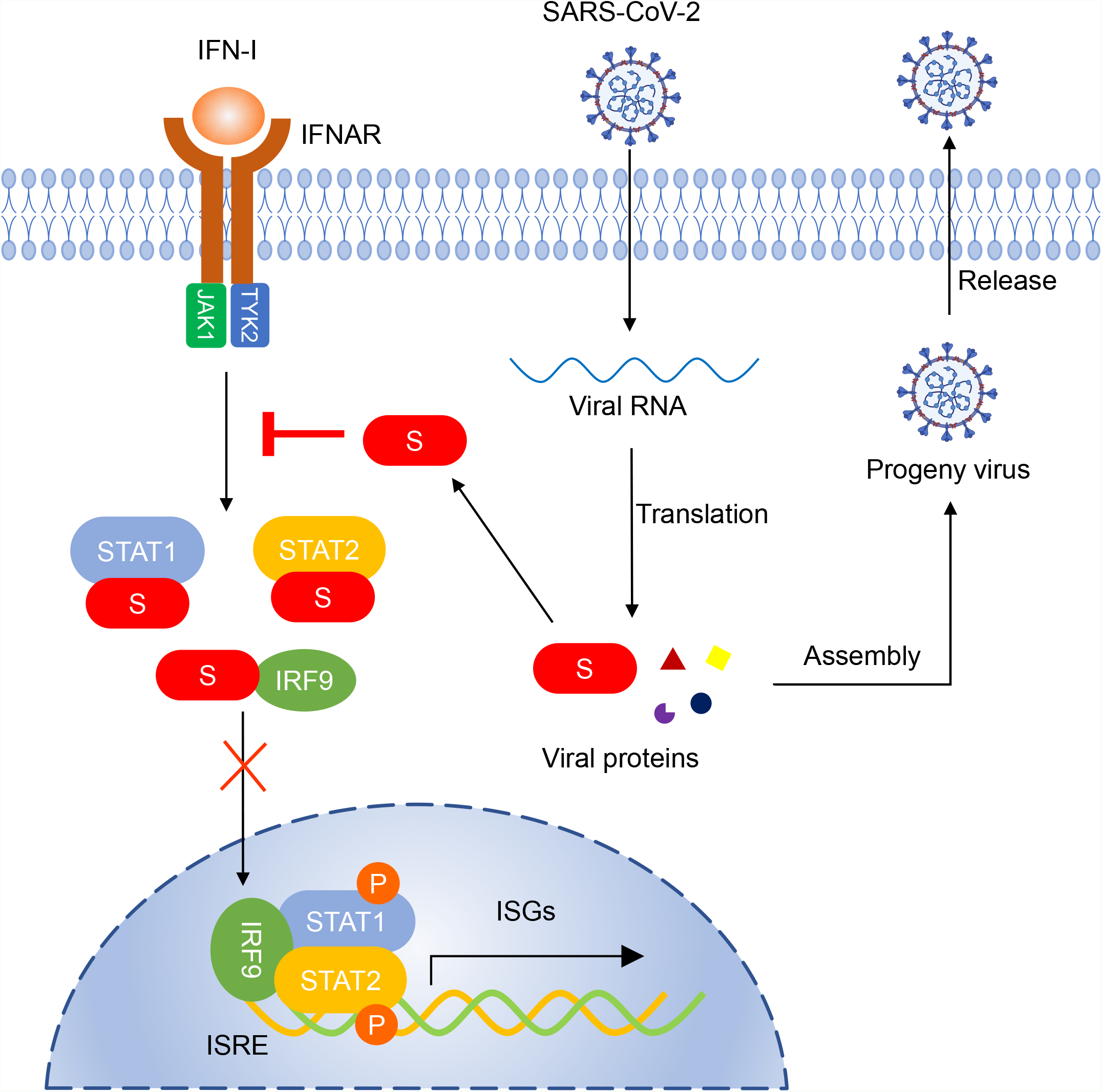
Working model. SARS-CoV-2 S protein interacts with STAT1/STAT2 and IRF9, suppresses the phosphorylation of STAT1 and STAT2, and hijacks STAT1 nuclear translocation, which blocks IFN-I signaling.

## Discussion

IFN response is the first line of defense against invading viruses during viral infection. Previous studies have shown that severe attenuation of IFN downstream cytokine expression is observed in patients with severe symptoms of SARS-CoV-2 infection (*24-26*). For example, IFN-I and ISG56 are almost not induced in the early stage of virus-infected cells but surged in the late time point. This delayed antiviral response may provide a window of activation and evaluation of IFN-I responses by SARS-CoV-2 (*2, 15, 27*). Our results indicate that the S protein antagonizes the IFN-I downstream JAK-STAT pathway (Fig. 6), partially explaining the attenuation of the IFN-I response in patients with COVID-19.

Increasing studies have suggested that SARS-CoV-2 may develop various strategies to limit effective IFN production, NSP1, NSP3, NSP6, NSP8, NSP12, NSP13, NSP14, ORF3a, ORF6, ORF7a, ORF7b, ORF8, ORF9b, M, and N proteins have been previously reported to inhibit both IFN-I production and downstream signaling (*15-19*). For example, ORF6 antagonizes IFN-I response via its C terminus and inhibits STAT1 nuclear translocation but not phosphorylation (*15*); another study reported that ORF6 hijacks Nup98 to block STAT nuclear import (*18*). M had proved to affect the RIG-I and TRIM25 to inhibit NF-κB signaling, impairing the phosphorylation of STAT1 and the nuclear translocation of STAT1 (*28*). N suppressed phosphorylation and the nuclear translocation of STAT1 and STAT2 (*17*). However, those inhibitory mechanisms remain for further study. Here, we reported S protein could target the JAK-STAT pathway to block the production of downstream ISGs, then inhibit the host antiviral immune signaling pathway. In subsequent studies, we will further focus on the synergistic effects of these proteins in evading the host’s innate immunity.

SARS-CoV-2 S protein is composed of two subunits, S1 and S2, playing different roles in viral infection. S1 contains the receptor-binding domain that binds to the receptor on the host cell (*29, 30*); S2 contains the fusion peptide required for viral entry, making it one of the most important targets for drug design (*31, 32*). In our data, the S1 subunit (NTD, RBD, SD1/2) and S2 subunit of S protein might play different functions. NTD showed no inhibitory effect, RBD and SD1/2 only slightly inhibited the activity of ISRE. The S2 domain plays the most critical inhibitory role. The S2 domain is the most conserved in SARS-CoV-2 variants, while the mutation sites that affect infectivity and neutralization escape are almost all concentrated in the S1 subunit (*33, 34*). Based on this, we speculate that S protein inhibition of the JAK-STAT pathway is not affected by the evolution of SARS-CoV-2. Therefore, we tested the effect of the S protein on ISRE of several Variants of Concern (VOC). As expected, our data showed that all of them significantly inhibited the activity of ISRE without significant difference compared with wild-type strains.

Moreover, our analysis shows that the S2 domain had high similarities in different human coronaviruses. So, we further tested whether the S protein of other human coronaviruses, SARS-CoV, MERS-CoV, HCoV-229E, HCoV-NL63, and HCoV-HKU-1, had the same effect. To our surprise, the inhibition of the JAK-STAT pathway by the S protein of coronavirus seemed to be conservative. Still, the underlying mechanism in other human coronaviruses remains for further study. Therefore, this study provides a new direction for finding common therapeutic targets of coronaviruses.

In summary, this study is the first to investigate the role of SARS-CoV-2 S protein in resisting IFN-I mediated innate antiviral immunity. Our study expands the understanding of SARS-CoV-2 and other human coronaviruses in evading antiviral immunity strategies, thus providing a theoretical basis for human anti-coronavirus immunity and understanding the interaction between host and coronavirus.

## Materials and Methods

### Cell Culture

293T and HeLa cells were obtained from American Type Culture Collection (ATCC) and maintained in Dulbecco’s modified Eagle medium (DMEM, Gibco) supplemented with 10% fetal bovine serum (FBS) and 1% penicillin & streptomycin (P/S, Gibco) at 37°C in an incubator with 5% CO_2_.

### Animal and Ethics Statement

Heterozygous B6/JGpt-H11em1Cin(K18-ACE2) /Gpt mice (K18-hACE2 KI mice) were purchased from GemPharmatech Co., Ltd. (Nanjing, China). C57BL/6 mice were purchased from Beijing Vital River Laboratory Animal Technology Co. Ltd. (Beijing, China). Mice were maintained and bred in individually ventilated cages (IVC) in a specific pathogen-free (SPF) environment. Animal experiments were executed by certified staff in the Center for Animal Experiments of Wuhan University (AAALAC #001274), approved by the Institutional Animal Care and Use Committee (AUP #WP2020-0819). And the protocols and procedures of infectious SARS-CoV-2 virus under Animal Biosafety Level-III Laboratory facility were approved by the Institutional Biosafety Committee (IBC, Protocol #S01320080E). All samples’ inactivation was performed according to IBC approved standard procedures for removal of specimens from high containment.

### Mice Infection and Sample Processing

For the mouse experiments, 12-18-week-old K18-hACE2 KI mice were anesthetized with isoflurane and intranasally inoculated with 50 μL of DMEM containing 2.5×10^2^ or 3.0×10^5^ PFU of SARS-CoV-2, respectively. K18-hACE2 KI mice intranasally inoculated with an equal volume of DMEM were considered as a mock control. Mice (at least 3 mice per group) were euthanized at 2, 4, and 6 dpi to collect lung tissue samples, which were then weighed, homogenized in PBS, and clarified by centrifugation at 5,000 rpm for 5 min. The supernatants were aliquoted and stored at -80°C before use.

### Quantitative Real-time PCR

The mRNA levels of the indicated genes were quantified via quantitative PCR with reverse transcription (qRT-PCR). Total RNA was extracted by using TRIzol (Invitrogen™,15596018) and reverse transcript to cDNA by PrimeScrip RT reagent Kit (Takara cat #RR037A). The corresponding cDNAs were quantified by using Hieff qPCR SYBR Green Master Mix (Yeason cat #11202ES03). The Primer sequences were provided below. All mRNA levels were normalized to the β-actin level or mActb level.

The primers were listed below:

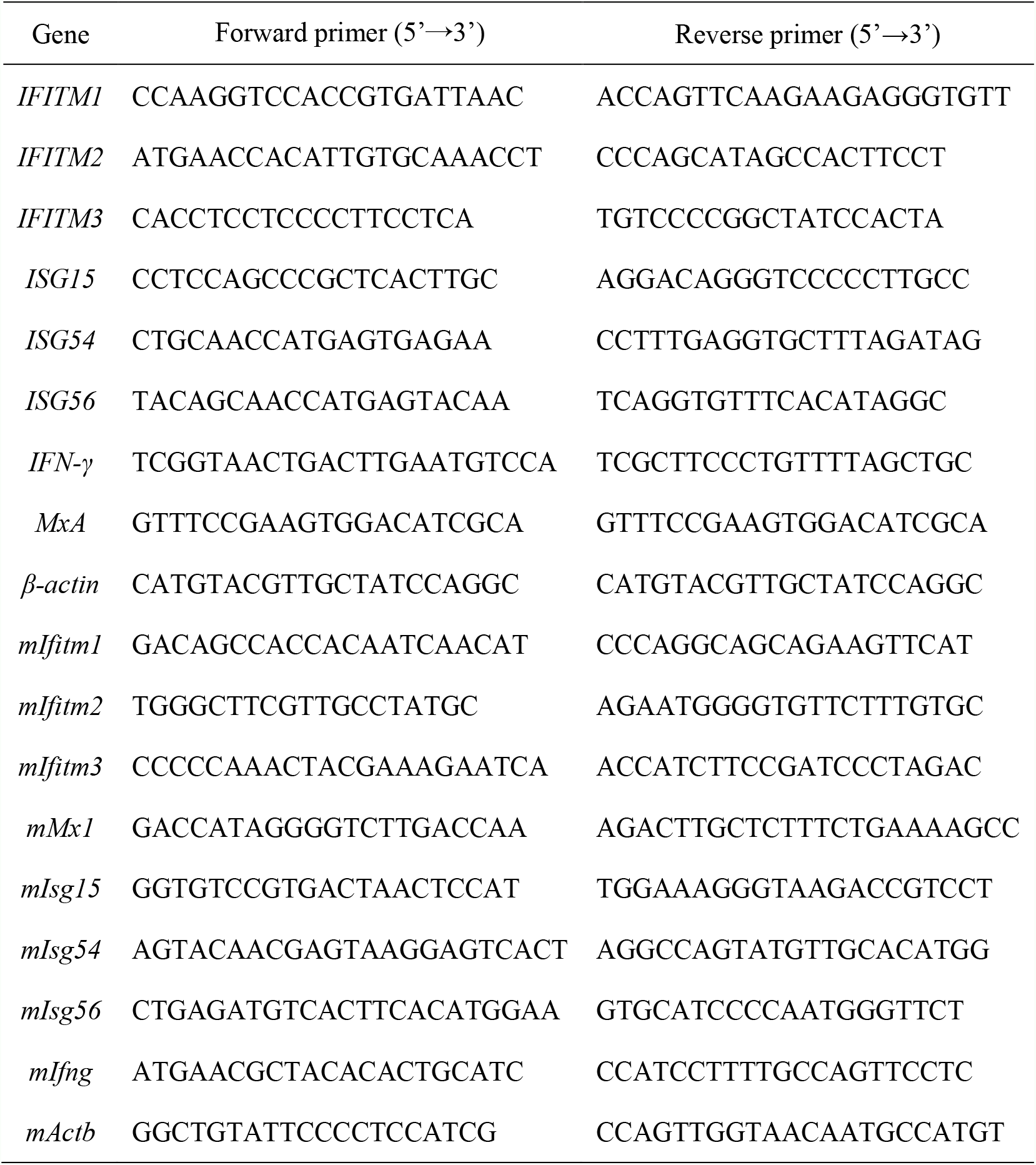

### Plasmid and Transfection

The plasmid encoding SARS-CoV-2 ORFs were cloned into pCAGGS vector. The plasmid encoding SARS-CoV-2 S protein with Flag-tag was purchased from SinoBiological (Cat. VG40589-NF). Mammalian expression plasmids for HA-, Flag-, or myc-tagged STAT1, STAT2, and IRF9 were constructed by standard molecular biology techniques. pISRE-Luc and pRL-SV40 were preserved in our laboratory. The S protein of SARS-CoV-2 variants (Alpha, Beta, Gamma, and Delta), SARS-CoV, MERS-CoV, 229E, NL63, HKU-1, and NTD, RBD, SD1&SD2, S2 of SARS-CoV-2 S protein was cloned into pCAGGS vector. The DNA transfection reagent using Lipofectamine 2000 (Invitrogen) and conducted according to the manufacturer’s protocol.

### Luciferase Assay

293T cells were seeded in 24-well plates and were transfected with a control plasmid or plasmid expressing indicated gene along with ISRE-, IFITM3-, or MxA-promoter firefly luciferase reporter plasmid and pRL-SV40 for 24 h. Subsequently, cells were stimulated with IFN-I (1000 U/mL) for 8 h, then cells were harvested and cell lysates were used to determine luciferase using a Dual-Luciferase Reporter Assay System (Promega). The firefly luciferase activities were normalized to Renilla luciferase activities.

### Western Blot

Cells were lysed in RIPA buffer on ice for 30 min, separated by using SDS-PAGE, and subjected to Western blot analysis. Mouse monoclonal HA-tag antibody (Biolegend, 901515, 1:1000), mouse monoclonal Flag-tag antibody (Sigma, F3165, 1:1000), rabbit monoclonal HA-tag antibody (CST, 3724, 1:1000), rabbit monoclonal Flag-tag antibody (CST, 14793, 1:1000), rabbit monoclonal myc-tag antibody (CST, 2278, 1:1000), rabbit polyclonal SARS-CoV-2 Spike antibody (SinoBiological, 40591-T62, 1:5000) and rabbit polyclonal anti-tubulin antibody (Antgene, ANT327, 1:2000) were purchased commercially. Peroxidase conjugated secondary antibodies (Antgene, 1: 5000) were applied accordingly, followed by image development with a Chemiluminescent HRP Substrate Kit (Millipore Corporation).

### Co-immunoprecipitation (Co-IP)

Cells were harvested and washed with ice-cold PBS and lysed in RIPA buffer and protease inhibitor (Roche, Cat.04693116001) for 30min on ice. The lysates were incubated with anti-HA or anti-Flag Agarose beads (Sigma, A2095, and A9469) as indicated overnight at 4°C. The immunoprecipitations were separated by SDS-PAGE and analyzed by immunoblotting.

### Immunofluorescence Microscopy

HeLa cells grown on glass coverslips were fixed in 4% paraformaldehyde (in PBS) for 15 min at room temperature and permeabilized with 0.1% Triton X-100 (in PBS) for 15min at room temperature. Subsequently, the cells were blocked with 1% BSA (in PBS) for 1 h and incubated with the rabbit monoclonal anti-HA antibody (CST, 3724, 1:500) or mouse monoclonal anti-Flag antibody (Sigma, F3165, 1:100) overnight at 4°C. Cells were then washed with PBS and stained with secondary antibodies (Alexa Fluor R488, Invitrogen; Alexa Fluor M568, Invitrogen) for 45 min at room temperature in the dark and then washed three times with PBS. The cell nucleus was stained with DAPI (Sigma, D9542) according to the standard protocols. Cell imaging was performed on a Zeiss LSM880 with an Airyscan confocal laser scanning microscope.

### Homology Analysis

The full-length Spike protein sequences of SARS-CoV-2, SARS-CoV, MERS-CoV, 229E, NL63, and HKU-1 were aligned using MUSCLE. The aligned sequences were further confirmed using similarity plot implemented in Simplot 3.5.1.

### Statistical Analysis

Data were analyzed with GraphPad Prism 8 software. Data are expressed as the mean ± standard deviation (SD). Comparisons of groups were performed using two tailed Student’s t-test or One-Way ANOVA test. The values *p < 0.05, **p < 0.01, ***p < 0.001, and ****p < 0.0001 were considered significant.

## Acknowledgments

This work was supported by the National Natural Science Foundation of China (grants 31922004 to K.X.), the National Key R&D Program (2021YFC2300702 to L.Z.), the Innovation Team Research Program of Hubei Province (2020CFA015 to K.L. and K.X.), and the Innovation Team Research Program of Fundamental Research Funds for the Central Universities (2042022kf1188 to K.X.). We are grateful to a special fund for COVID-19 Research of Wuhan University, the fund from Taikang Insurance Group Co., Ltd and Beijing Taikang Yicai Foundation, and the Fundamental Research Funds for the Central Universities for their great support of this work.

## Author Contributions

K.X. conceived the project. K.X. and L.Z. designed the experiments. W.N., W.L., and Z.C. conducted the luciferase assay, the IFN treatment experiments, the Co-IP experiments, the immunofluorescence study, and data analysis. W.N., W.L., J.Y., X.G., and D.Z. constructed the plasmids of SARS-CoV-2 ORF proteins. W.G. and Z.W. performed the gene mutagenesis and protein purification assays. Y.Z. and S.L. prepared the spike variant plasmids and cells. L.Z. and Y.Z. performed live SARS-CoV-2 study and animal infection experiments. X.W., C.Z., M.T., K.L., and Y.C. provided technical supports and materials. K.X., L.Z., W.N., W.L., and Z.C. wrote the manuscript with input from all the other authors. All authors read and approved the final manuscript.

